# A data-driven model of macrophage polarization states reveals an IFN macrophage signature in active Crohn’s disease

**DOI:** 10.1101/2025.09.24.678388

**Authors:** Sarina C. Lowe, Madelaine Leitman, Noah Yan, Alexander Hoffmann

**Author notes:** shared first authorship.

## Abstract

Macrophages are key innate immune cells responsible for initiating and coordinating immune responses. A major determinant of macrophage function is their tissue-context-dependent polarization state, regulated by the cytokines present in the tissue microenvironment. Yet, the capability of characterizing macrophage polarization states in clinical studies remains limited. Here, we used a defined set of cytokines to polarize human PBMC derived macrophages and determine their transcriptional signatures and stimulus responsiveness. The resultant atlas of transcriptional signatures for human macrophage polarization states was applied to a dataset of intestinal biopsies from Crohn’s disease patients and healthy controls. Our analysis identified a dominant population of IFNγ-polarized macrophages in areas of active Crohn’s intestinal inflammation and a loss of wound healing IL-4-, IL-10- and IL-13-polarized macrophages. This study demonstrates that *in vitro* datasets of macrophages in defined conditions can be leveraged to interpret the functionality of cells transcriptional profiled in clinical studies.

## Introduction

Macrophages serve diverse roles within the immune system, ranging from the acute response to pathogens to wound healing and maintenance of homeostasis. Tissue microenvironmental cytokines may cause macrophage polarization resulting in changes in their functional repertoire. These adaptations affect macrophage morphology, signaling dynamics, gene expression, and cytokine production (1–7). This is particularly relevant in the intestine, where most macrophages originate from circulating monocytes (8) that extravasate into the microenvironment of the intestinal lamina propria, where they have a lifespan of 3-5 weeks (9).

In Crohn’s disease, macrophages are preferentially enriched at sites of active inflammation and are a key source of proinflammatory cytokines that contribute to disease pathogenesis (4,10). Although many of the macrophage-derived cytokines (i.e. TNF, IL-12, IL-23) are therapeutic targets, the repertoire of macrophage polarization states that drive cytokine production in Crohn’s disease is unknown.

Single-cell RNA sequencing (scRNA-seq) enables multi-dimensional interrogation of cell states. Applied to *in vitro*-cultured macrophages exposed to specific cytokines, it reveals transcriptional profiles that define polarization states. This reference atlas can then be leveraged to analyze *in vivo* patient-derived scRNA-seq data, enabling characterization of macrophage populations in Crohn’s disease. Prior studies of macrophage subtypes in Crohn’s disease have been limited to categorization as ‘inflammatory’ and ‘noninflammatory’ based on biomarker expression, or gene expression clusters with little clinical or biologic significance (11). Here we produced an extensive scRNA-seq dataset of *in-vitro*-polarized macrophages that we leveraged to interpret *in vivo* scRN-seq data produced from Crohn’s disease clinical samples. As a result, we identified distinct macrophage subsets that are more or less abundant in active Crohn’s disease lesions and regions of healing.

## Methods

### In vitro macrophage culture, polarization, and stimulation

Human blood from a deidentified donor was obtained from the UCLA CFAR Centralized Laboratory Support Core. Peripheral blood mononuclear cells (PBMCs) were isolated from blood by Ficoll density centrifugation followed by CD14+ bead selection for monocytes. The monocytes were then plated on 6-well tissue culture plates at a density of 5 x 10^5^ cells/well, and cultured in RPMI media supplemented with 10% ES fetal bovine serum and 20 ng/ml human macrophage colony-stimulating factor. Media was replaced 4 days after isolation, and after 6 days the cells were polarized for 24 hours with either 100 U/mL IFN-β (PBL Assay Science, 11415-1), 20 ng/mL IFN-γ (Peprotech, 300-02), 50 ng/mL IL-10 (R&D, 217-IL), 40 ng/mL IL-13 (R&D, 213-ILB), 20 ng/mL IL-4 (R&D, 204-IL), or left naïve. After 24 hours of polarization, the cells were stimulated for 2 hours with LPS 10 ng/ml (Sigma-Aldrich, B5:055), TNF 100 μg/ml (R&D, 210-TA), or left unstimulated.

### In-vitro-polarized macrophage scRNA-seq and analysis

Cells were lifted and the samples were multiplexed using the BD Human Single-Cell Multiplexing Kit (BD, 633781) and loaded onto the cartridge following the manufacturer’s instructions (BD Rhapsody #210967). Targeted mRNA amplification was performed utilizing the Immune Response Panel Hs primer panel (BD, 633750), with subsequent library preparation. The pooled library was sequenced with paired-end 100 bp reads on an Illumina NovaSeq X Plus. The raw FASTQ files and reference for the Immune Response Panel were run through the BD Rhapsody Sequence Analysis Pipeline which includes quality filtering with read overlap detection, trimming and filtering, followed by identification of the cell barcode and unique molecular identifier (UMI), alignment to the human genome, and sample determination from the multiplexed sample tags, resulting in an output of adjusted molecule counts.

### In vivo terminal ileum scRNA-seq data processing and annotation

Published single-cell RNA sequencing (scRNA-seq) data was obtained from terminal ileal (TI) biopsies of Crohn’s disease patients (12). Immune cells were separated from stromal and epithelial cells based on annotations from the original study (12) and prepared for downstream analysis with log-normalization and standardization of expression. A UMAP was constructed using principal component analysis (PCA) reduction, and cell types were labeled by manual annotation of the top 20 differentially expressed genes for each cluster of immune cells. Macrophages were identified by high expression of canonical macrophage genes (including C1QA, C1QB, C1QC, CD14, and FCN1). The immune cell clusters were relabeled by manual annotation, and a macrophage-only object was subsequently processed independently.

### Integration of in vivo and in vitro macrophage data

To integrate the *in vivo* TI macrophage data composed of 28,923 genes with the *in-vitro*-polarized macrophage data with 344 genes, both datasets were standardized by retaining only the 325 overlapping genes for downstream analysis (13). With the *in vivo* TI macrophage dataset as the reference, Seurat’s anchor-based integration framework was used with the *in-vitro*-polarized dataset as the query. The *in-vitro*-polarized macrophages were assigned to an *in vivo* TI unsupervised cluster (1, 2, or 3), mapping the *in-vitro*-polarized cells’ PCA profile onto the reference *in vivo* TI macrophage’s PCA space. The most transcriptionally similar *in vivo* TI cluster based on a mutual nearest-neighbor classification (14) (k=30) was assigned to each *in-vitro*-polarized macrophage cell profile using the highest similarity score.

## Results

We developed a workflow to better characterize the macrophage cell states found in Crohn’s disease (Fig.1). We first systematically profiled *in-vitro*-polarized human macrophages exposed to defined cytokine polarization conditions and identified polarization gene signatures. Applying these to clinical samples enabled identification of specific intestinal macrophage subpopulations in Crohn’s patients.

**Figure 1.**
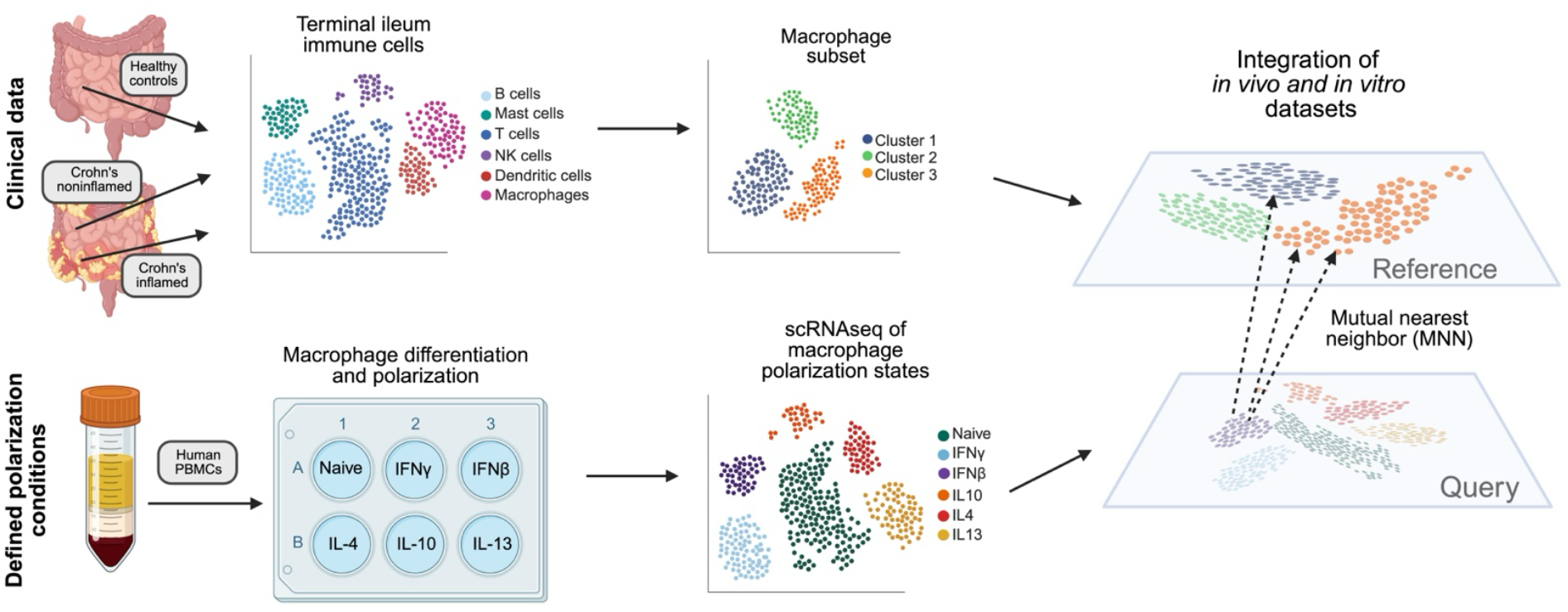
Overview of an experimental and analytic workflow to better characterze macrophage states in clinical scRNA-seq datasets. Workflow used to generate an *in vitro* atlas of macrophage polarization states and apply it to functionally annotate an *in vivo* Crohn’s clinical dataset. Peripheral blood mononuclear cells (PBMCs) were isolated from human blood and differentiated into macrophages. After six days, cells were polarized with IFNγ, IFNβ, IL-4, IL-10 or IL-13, left unstimulated or stimulated with LPS or TNF, and processed for scRNA-seq using the BD Rhapsody platform. *In vivo* intestinal immune cell data from healthy controls and Crohn’s disease patients (Kong et al (12)) were re-analyzed, and macrophages were identified by annotation of cluster-specific differentially expressed genes. Using 325 shared genes and a mutual nearest neighbors approach to identify anchors, the *in vivo* and *in vitro* datasets were integrated into a shared space, enabling characterization of the polarizating state of macrophages present in clinical samples.

To prolife single macrophage polarization states we cultured PBMC-derived macrophages and polarized them with individual cytokines from a larger panel. We used an established sc-RNAseq targeted gene platform with a select panel of immune response genes that show stimulus-specific expression and assess the functional state of macrophages (15). The *in-vitro*-polarized macrophages showed distinct transcriptional profiles after 24 hours of polarization with IFNγ, IFNβ, IL-4, IL-10 and IL-13 (Fig. 2A and 2B).

**Figure 2.**
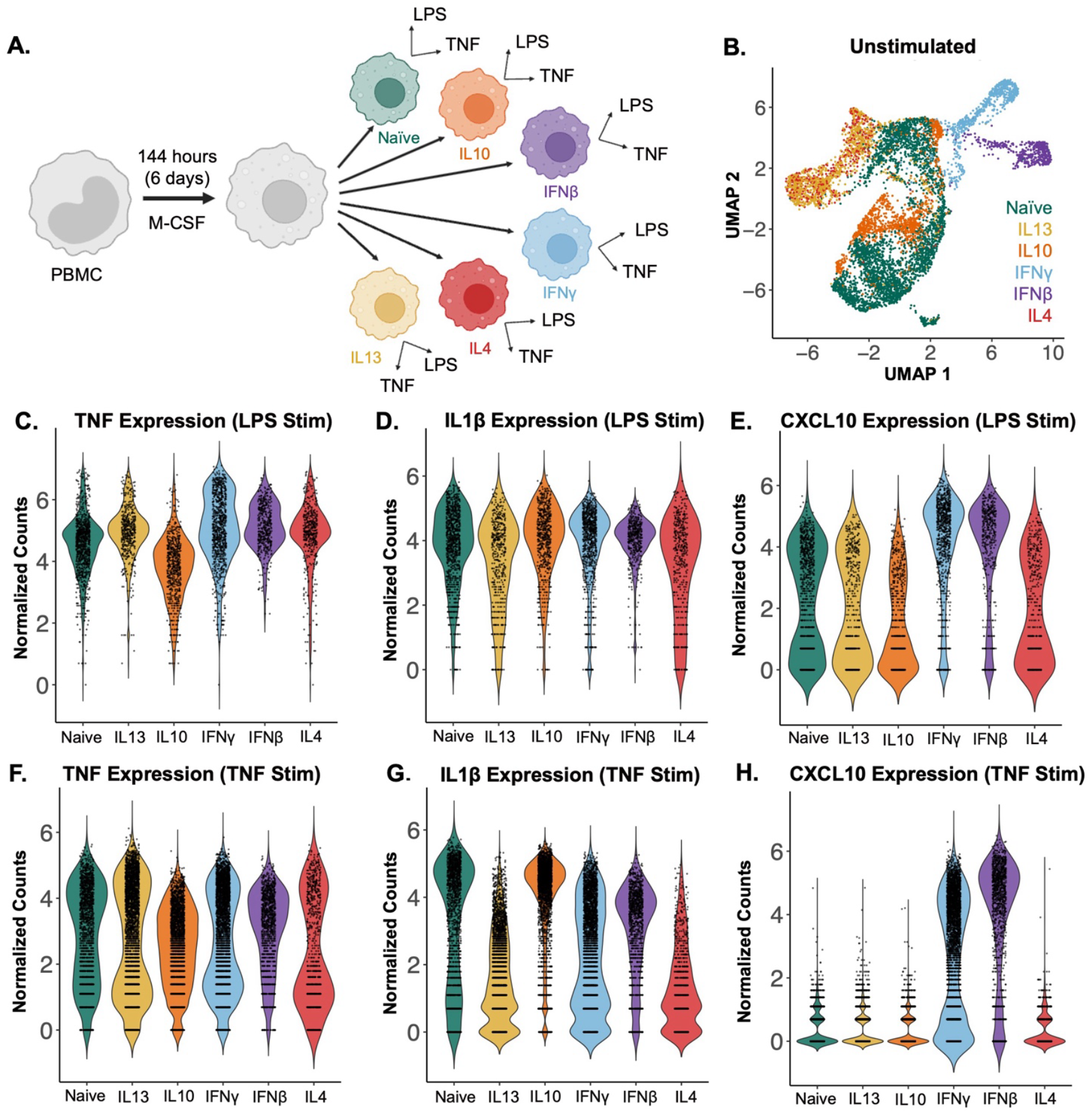
*In vitro* polarization and stimulation of human PBMC-derived macrophages define an atlas of macrophage polarization states. (A) Experimental overview: monocytes were isolated from PBMCs by CD14-positive selection and differentiated in M-CSF. After six days the polarizing cytokines (IL-4, IL-10, IL-13, IFNγ or IFNβ) were applied for 24 hours. Polarized macrophages were then stimulated with either LPS (10ng/ml) or TNF (100ug/ml) for 2 hours prior to RNA isolation and sequencing. (B) UMAP of PBMC-derived day 6 macrophages polarized for 24 hours without acute stimulation. The samples are colored by polarization condition with naïve in green, IL-13 in yellow, IL-10 in orange, IFNγ in blue, IFNβ in purple and IL-4 in red. (C-E) Violin plots of TNF, IL1β and CXCL10 expression in polarized macrophages stimulated with LPS. (F-H) Violin plots of TNF, IL1β and CXCL10 expression in polarized macrophages stimulated with TNF. Each dot represents an individual cell while the density curve depicts the distribution of numeric data. Counts were normalized by regularized negative binomial regression.

To assess functional differences between these polarization states, the macrophages were subsequently stimulated with either lipopolysaccharide (LPS) or tumor necrosis factor (TNF). The stimulated macrophages showed differences in expression of key inflammatory cytokines and chemokines when normalized by regularized negative binomial regression. In response to LPS stimulation, IFNγ- and IFNβ-polarized macrophages showed increased TNF expression compared to naïve cells, while the IL-10-polarized macrophages displayed reduced TNF expression (Fig. 2C).

However, this polarization effect on LPS-responsive gene expression was not uniform across all cytokines and chemokines. IL1β expression showed minimal variation between polarization conditions (Fig. 2D), whereas CXCL10 expression differed markedly, with IFNγ- and IFNβ-polarized macrophages exhibiting much higher levels of expression than other conditions (Fig. 2E). Notably, these gene-specific polarization effects differed between LPS and TNF stimulation. With TNF stimulation, there was little difference in subsequent TNF expression between polarization conditions (Fig. 2F), but an increase in IL1β expression in the naïve and IL-10 polarized macrophages compared to the other conditions (Fig. 2G), while CXCL10 was not expressed in any condition but the IFNγ and IFNβ polarized (Fig. 2H). By assessing inflammatory gene expression following stimulation, we identified functional differences in macrophages across polarization conditions.

After establishing an atlas of polarized macrophage transcriptional states, we leveraged it to better characterize the macrophage populations present in Crohn’s disease. Terminal ileum data was available from 27 Crohn’s patients and 9 healthy donors (Figs. 3A and 3B), with a higher percentage of macrophages in the Crohn’s samples, predominantly in areas of active inflammation (Fig. 3C).

**Figure 3.**
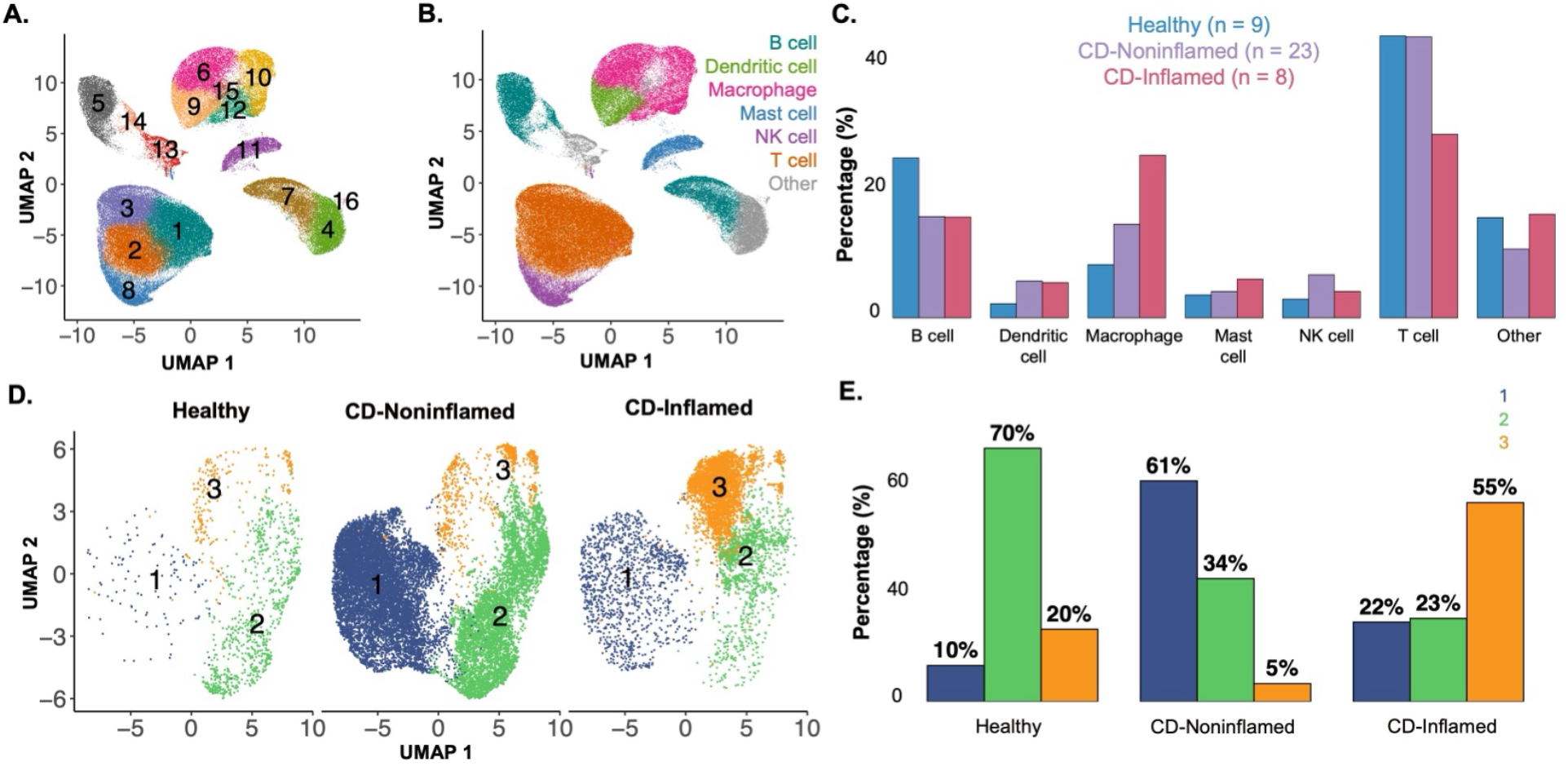
Immune cell compositions of terminal ileal samples using established marker genes. (A) UMAP of terminal ileum immune cells from healthy donors (N=9) and Crohn’s patients (N=27), from both noninflamed and inflamed regions. Immune cells were separated from stromal and epithelial cells based on annotations from the original study (Kong et al) and clustered in an unsupervised manner. The numbers delineate each individual cluster. (B) Immune cell types manually annotated based on the top 20 differentially expressed genes for each unsupervised cluster, with some clusters consolidated into broader cell type categories. The macrophages are shown in pink. (C) Proportion of immune cell types stratified by donor types (healthy in blue, Crohn’s noninflamed in purple, and Crohn’s inflamed in pink). (D) Unsupervised clustering of the macrophage subset, separated by donor type. Cluster 1 is marked in blue, cluster 2 in green, and cluster 3 in yellow. (E) Distribution of macrophages assigned to each of the three clusters (colors as in panel D), separated by donor type.

Macrophage-specific data from healthy donors (N=9), Crohn’s noninflamed biopsies (N=23), and Crohn’s inflamed biopsies (N=8) (Fig. 3D) revealed notable differences in macrophage subgroup distribution. Cluster 2 macrophages were dominant in biopsies from healthy donors, while cluster 1 macrophages were dominant in Crohn’s noninflamed biopsies, and cluster 3 in Crohn’s inflamed biopsies (Fig. 3E).

The *in-vitro*-polarized macrophage data (Fig. 4A) was integrated with the *in vivo* Crohn’s disease terminal ileum dataset (Fig. 4B) using a shared set of 325 genes. Integration was performed using Seurat’s SCTransform-based normalization to mitigate technical noise. Each cell was then projected into a common space by applying an anchor-derived transformation (16). This anchor strategy identifies pairs of transcriptionally similar cells across the datasets, allowing the two datasets to be aligned in a common low-dimensional space. After integration, the polarization labels from the *in vitro* macrophages were projected onto the *in vivo* dataset using a weighted nearest-neighbor (k=30) classification (Fig. 4C), where each *in vitro* macrophage was assigned the label of its most transcriptionally similar *in vivo* cell based on the highest similarity score (Fig. 4D).

**Figure 4.**
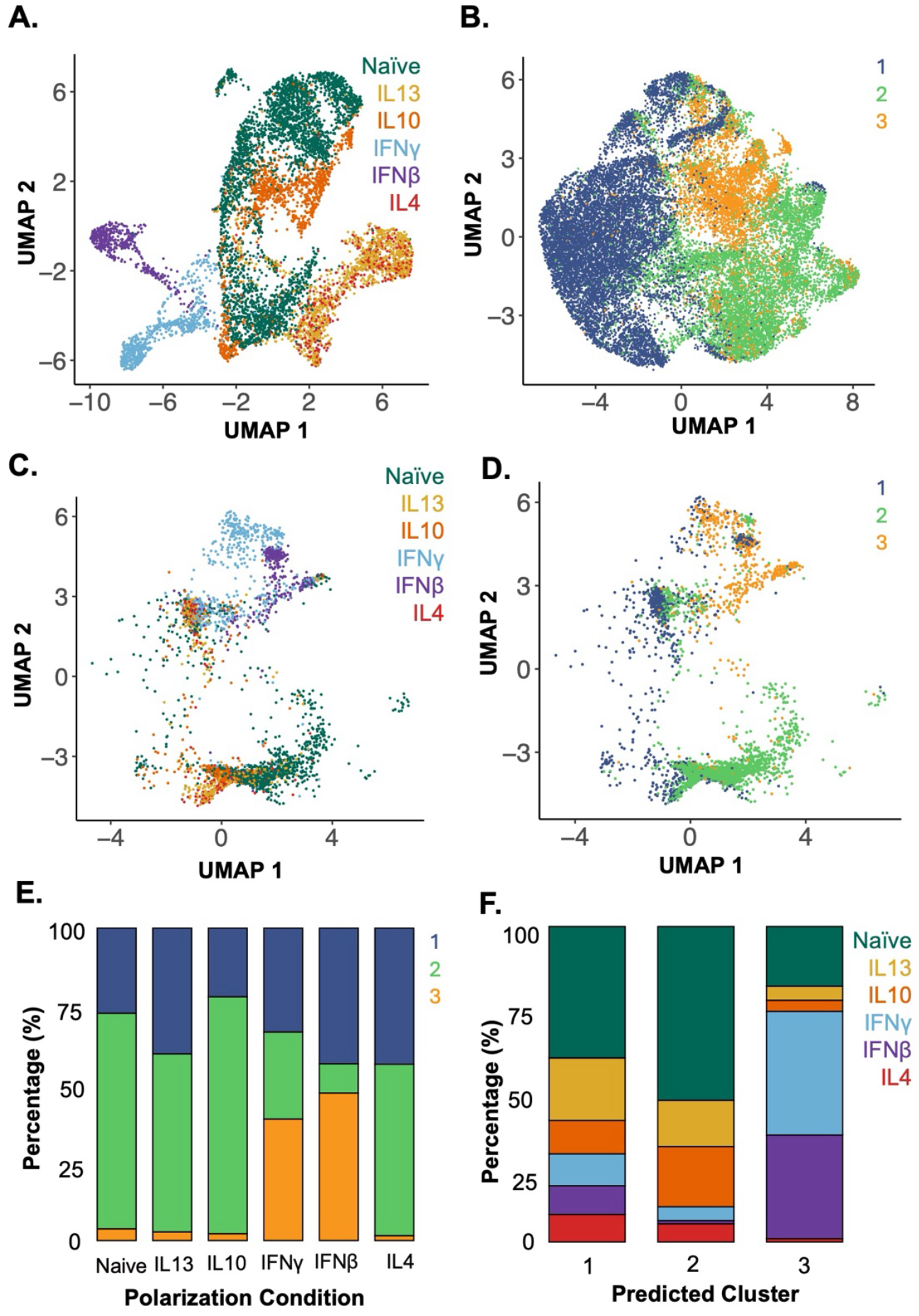
Applying a model trained on *in-vitro-*polarized macrophage datasets identifies the polarization states of *in vivo* terminal ileal macrophages in clinical scRNA-seq data. (A) UMAP of i*n-vitro*-polarized macrophage scRNA-seq data using the shared set of 325 genes (colors as in Fig. 2). (B) UMAP of *in vivo* terminal ileal macrophages labeled by assigned cluster (colors as in Fig. 3) after reducing the 28,923 gene dataset to the 325 genes shared with the *in vitro* atlas. (C) *In-vitro*-polarized macrophages integrated with the terminal ileal dataset using SCTransform anchors and mapped onto a shared UMAP, labeled by polarization condition. (D) Assignment of *in-vitro-*polarized macrophages to *in vivo* clusters 1-3 using a weighted mutual nearest neighbor approach. (E) Proportion of cells from each polarization condition assigned to each *in vivo* macrophage cluster by mutual nearest-neighbor classification. (F) *In vivo* cluster assignments made by mutual nearest-neighbor classification displayed by polarization condition.

When examining the different macrophage polarization states and their assigned *in vivo* clusters, the majority of naïve, IL-4, IL-10 and IL-13 *in-vitro*-polarized macrophages aligned to cluster 2, the cluster most highly represented in biopsies from healthy donors. IFNβ- and IFNγ-polarized macrophages aligned with cluster 3, which best represented macrophages from areas of active Crohn’s disease (Fig. 4E). This integrated data and cluster assignments were used to examine the polarization states of the *in vivo* terminal ileum macrophage populations (Fig. 4F). Macrophages in cluster 1 were distributed throughout the range of polarization conditions with the largest portion of cells aligning to the naïve and IL-13-polarized states. Cluster 2 was enriched in cells aligning to the naïve condition and under-enriched in IFN-polarized cells. Lastly, in cluster 3, which was highly enriched in macrophages from inflamed Crohn’s subjects, the majority of macrophages aligned to the IFNβ- and IFNγ-polarized conditions (17). These analyses were able to characterize the polarization state of macrophages within biopsies based on an atlas of *in-vitro*-defined polarized macrophages.

## Discussion

Culturing macrophages with defined cytokines produced distinct populations of polarized macrophages. Not only did the different macrophage polarization states exhibit varying baseline transcriptomic profiles, but their gene expression in response to stimuli, both a pathogen associated molecular pattern (PAMP), lipopolysaccharide (LPS), and a proinflammatory cytokine, tumor necrosis factor (TNF), differed. In particular, the expression of inflammatory mediators such as cytokines and chemokines varied between polarization state and stimuli condition, highlighting the impact of polarization on macrophage function.

By applying a model based on *in vitro* single-cell macrophage polarization data to *in vivo* single-cell data from Crohn’s patients, this analysis revealed macrophages from inflamed Crohn’s samples have a transcriptional program closest to proinflammatory IFN-polarized macrophages at the expense of anti-inflammatory IL-4-, IL-10- and IL-13-polarized macrophages. The results support the notion of the cytokine microenvironment as a driver for distinct macrophage subsets in Crohn’s disease. IFNγ is largely produced by innate lymphoid cells (ILCs), NK cells, and activated T cells, and is elevated in the mucosa of patients with Crohn’s disease (18) along with animal models of colitis (19). From a macrophage standpoint, the IFNγ environment is significant because it is known to polarize towards a proinflammatory macrophage phenotype, which exhibits enhanced inflammatory responses with increased proinflammatory cytokine production (20). The development of IFNγ-polarized macrophages in response to an inflammatory environment further promotes intestinal inflammation as these macrophages are primed to produce more proinflammatory cytokines, activating and recruiting further immune effector cells.

In contrast, the cluster of macrophages from noninflamed segments (cluster 1) exhibited an even distribution of macrophage polarization states, and particularly more IL-4-, IL-10- and IL-13-polarized macrophages. These samples were from patients with Crohn’s disease, but with inactive disease, or no evidence of active inflammation in the biopsied portion of terminal ileum. The distribution of polarization states skewing towards more IL-4-, IL-10- and IL-13-polarized macrophages is consistent with the pivotal role of anti-inflammatory macrophages in wound healing and resolution of inflammation (4).

The findings highlight the pathogenic role of IFN-polarized macrophages in active Crohn’s intestinal inflammation, and the healing role of anti-inflammatory IL-4-, IL-10- and IL-13-polarized macrophages. This could impact therapy by identifying patients with a predominant interferon macrophage signature who may benefit from JAK/STAT inhibition, which is a therapeutic option for Crohn’s disease. Inhibition of this pathway could not only block acute inflammatory signaling downstream of IFNs but also prevent IFNγ-induced macrophage polarization. The presence of IL-4-, IL-10- and IL-13-polarized macrophages in biopsies from noninflamed/healed areas of Crohn’s disease also spotlights using anti-inflammatory cytokines to steer macrophages away from the pathogenic inflammatory state towards an anti-inflammatory polarization state as a viable therapeutic approach.

This work pioneers the generation of an *in vitro* defined macrophage polarization atlas characterized by transcriptomes and applying them to better characterize macrophage populations in Crohn’s disease. Prior investigations of macrophages in Crohn’s disease relied on markers of ‘inflammatory’ M1 or ‘anti-inflammatory’ M2 macrophages. The present study shows that single-cell RNA sequencing data of *in vitro* cytokine-specific polarization of macrophages can inform *in vivo* macrophage polarization states with potential clinical impact. The approach provides a robust characterization of macrophage states that identifies the key cytokines shaping the macrophage population in Crohn’s disease. These results highlight opportunities to possibly treat Crohn’s disease by altering the inflamed microenvironment, inhibiting the formation of pathogenic proinflammatory macrophages and shifting the population to immune suppressing anti-inflammatory macrophages. They also identify how rigorous *in vivo* characterization of cell types – beginning with transcriptomes and extending to cell type-specific functions – can advance our understanding of disease pathogenesis and inform treatment.

